# Selective targeting of oncogenic KRAS G12D using peptide nucleic acid oligomers attached to cell-penetrating peptides

**DOI:** 10.1101/2025.03.28.645837

**Authors:** Jayati Mondal, Dennis Lam, Mary E. Gerritsen, Tilmann M. Brotz, Jodi G. Kennedy, Bruce Rehlaender, Arthur J. Ross, Daniel E. Levy, Christopher A. Bonagura, William N. Lanzilotta, Frank McCormick, Jeffrey H. Rothman, Andrew L. Wolfe

## Abstract

KRAS is a proto-oncogene that contains activating mutations in up to 30% of tumors. Many conventional therapies inhibit both cancerous and normal cells, which may cause toxicity. Thus, programmable mutant-selective targeted inhibitors are needed. Peptide nucleic acids (PNAs) incorporate base sequences analogous to DNA, with modified peptide backbones instead of ribose-phosphate backbones, allowing PNAs to hybridize with DNA with high avidity to suppress transcription. Here, we developed KRAS G12D-selective PNA oligomers with novel cell-penetrating flanking regions. Fluorescein-labeled PNA oligomers displayed high uptake rates in cells and nuclei. Exposure to PNA-delivery peptide conjugates resulted in repression of KRAS G12D mRNA and protein expression within 2 hours and lasting up to 48 hours. Varying cell-penetrating peptide (CPP) compositions and lengths of complementary KRAS sequences were tested using dose-response cell viability assays. These experiments identified configurations that were effective at selectively preventing growth of on-target KRAS G12D cells, while relatively sparing off-target KRAS G12C cells. Electrophoretic mobility shift assays demonstrated *in vitro* binding and selectivity for KRAS G12D DNA sequences. CPP-PNA-G12D-1 was effective against a panel of pancreatic ductal adenocarcinoma cell lines and patient-derived xenografts *in vivo*. These results show promise for an enhanced PNA-delivery peptide conjugate strategy as both a tool for studying tumors driven by oncogenic point mutations and as a potential therapeutic strategy to selectively target mutant cancer cells.

## Introduction

Conventional chemotherapies non-selectively target growing tumor cells and normal cells causing high toxicities, highlighting the need for more selective and programmable targeted therapeutic strategies. KRAS (Kirsten Rat Sarcoma Viral Oncogene Homolog) is part of the RAS family of genes, which includes NRAS and HRAS, and plays a crucial role in driving proliferative cellular signaling pathways. KRAS proteins function as small GTPases that toggle between an active and inactive state, driven by the binding of guanosine triphosphate (GTP) or guanosine diphosphate (GDP). KRAS-GTP sends signals that regulate cell growth, division, and survival. KRAS mutations occur in 25–30% of all human cancers, including three of the worst prognosis cancers with approximately 20% of non-small cell lung cancers (NSCLCs), 40% of colorectal cancers (CRCs), and 73-90% of pancreatic ductal adenocarcinoma (PDAC).^1–3^ In normal cells, KRAS tightly regulates these signals, but mutations can cause KRAS to remain in a constitutively active state, promoting uncontrolled cell proliferation and cancerous growth. KRAS is most commonly mutated at codon 12.^4^ The only KRAS inhibitors currently approved by the FDA selectively target the G12C mutation in non-small cell lung cancer and colorectal cancer. These two compounds rely on irreversible covalent linkage between the inhibitor and the mutant cysteine, making them ineffective against other KRAS mutations.^5–7^ Experimental tri-complex pan-KRAS inhibitors currently in the clinic target other alleles, as well as wild-type KRAS, NRAS, HRAS, and other structurally related proteins.^5,8^ The extent to which pan-KRAS inhibitors’ ability to inhibit wild-type RAS proteins will result in undesirable toxicity in normal cells remains unclear, highlighting the need to develop alternative approaches to mutant KRAS inhibition.^1,8^

Peptide nucleic acids (PNAs) have nucleotide sequences analogous to DNA, except they have modified peptide backbones instead of ribose-phosphate backbones.^9^ PNAs were conceptualized and synthesized to create more stable, flexible molecules capable of resisting enzymatic degradation that bind strongly to DNA or RNA targets. Their design aimed to mimic the structure of DNA, but with a N-(2-aminoethyl)-glycine backbone linked to nucleobases instead of the typical sugar-phosphate structure of DNA and RNA.^10^ PNAs are capable of binding gene sequences ∼1000-fold more avidly than complementary native DNA. PNAs function to repress their targets by hybridizing with DNA at orders of magnitude higher affinity than the complementary DNA strand, strand-invading the duplex and blocking transcriptional machinery from transcribing the target sequence.^11,12^ PNAs represent a significant advancement in molecular biology and therapeutic research, especially in targeting of mutant genes and molecular diagnostics.^13^

Peptide nucleic acids are highly sequence-selective relative to alternatives such as small interfering RNAs, suggesting a therapeutic potential to selectively target cancerous cells with a specific point mutation using a complementary PNA.^14^ PNAs have been tested against mutant oncogenes including BRAF, and for suppression of oncogenic miRNAs and lncRNAs in cancer.^13,15–17^ Previous uses of KRAS G12D PNA employed lipofectamine delivery systems, liposomes, and/or cell-penetrating peptides to enter cells.^18,19^ Lipofectamine, a widely used transfection agent, forms complexes with PNAs, enhancing their entry into cells by temporarily destabilizing cell membranes. While effective in *in vitro* research, lipofectamine has limitations that prevent it from being viable in clinical settings.^18^

To optimize PNAs for specificity, we developed peptide nucleic acid oligomers conjugated to nuclear delivery peptides that can target and obstruct transcription, in a sequence- and allele-specific complementary manner against KRAS G12D, a target which is specific only to cancer cells. Due to the statistical challenge for the PNA conjugate finding its chromosomal target sequence, stabilizing the PNA oligomer against the chromosomal target would better ensure that the PNA conjugate remains within the chromosomal binding arena and increase the statistical likelihood of binding. Furthermore, as the termini are statistically the most configurationally labile portion of the cell-penetrating peptide/peptide nucleic acid (CPP-PNA) chain, its stabilization against the complementary target increases propensity for helical binding.^20^ The Zimm-Bragg helical model shows that an initial contact of two strands with helical structure forming propensity is the kinetically limiting event for helix formation.^20^ Stabilizing cationic termini, the most configurationally labile regions, towards the chromosome anionic ribose phosphates offers an effective route to maintaining contact reducing entropic obstacles towards helix formation (Fig. 1a). By better stabilizing these conjugates within the phospholipid bilayer of cellular membranes, the equilibrium of the CPP-PNA conjugate within the cell and nucleus is more quickly achieved as the membrane becomes less of a barrier.^16^ To achieve successful suppression of these regions, we extensively modified these PNA conjugates to improve their regional target stability and intracellular and nuclear delivery (Fig. 1a and Supplementary Fig. 1). We employed KRAS G12D and control cell lines to evaluate the ability of CPP-PNA conjugates targeting KRAS G12D to localize to nuclei, block KRAS G12D production, selectively impede cell viability of G12D cells, and inhibit G12D-driven tumor growth *in vivo*. These PNAs penetrated cells and repressed expression of mutant KRAS G12D. The PNA oligomer differentiated sequences differing by a single nucleotide base, and were effective at suppressing tumor growth in patient-derived xenografts with KRAS G12D. Our findings support the efficacy and the specificity of this improved CPP-PNA conjugate strategy.

**Figure 1.**
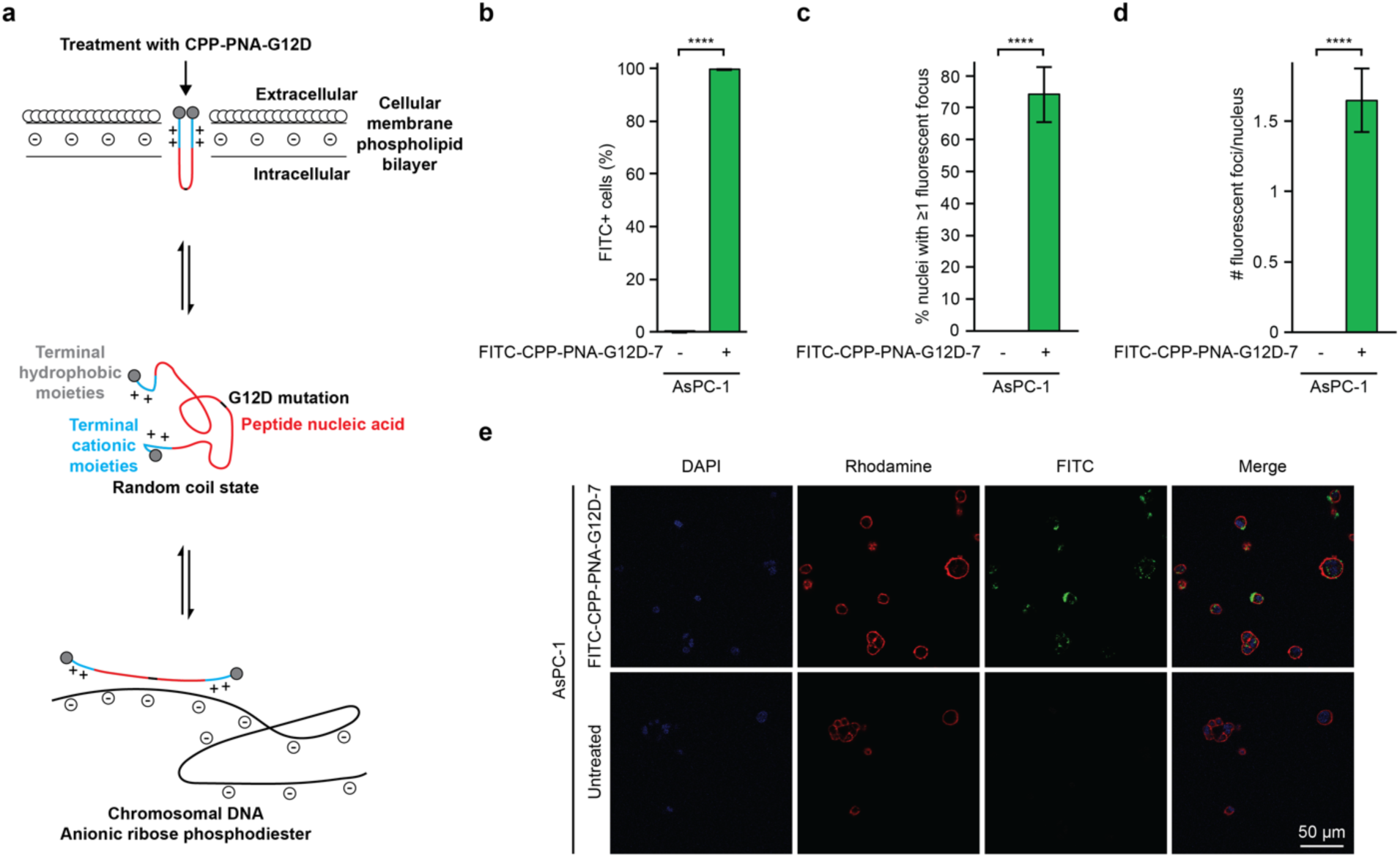
Cell-penetrating peptide PNA conjugates enter cells and nuclei. **a,** Model of CPP-PNA-G12D action. Terminal hydrophobic and cationic moieties are engaged during passage through membranes. CPP-PNA conjugates default to a random coil conformation. In the nucleus, CPP-PNA conjugates bind to complementary antisense genomic DNA. Targeting the antisense DNA strand precludes binding to mRNA. PNA-CPP-G12D binds directly to the residue 12 mutation site (thin black line) and the surrounding nucleotides, rather than the promoter regions targeted by some other PNAs, allowing for specificity to a mutation of choice. An accessible chromatin conformation is likely because KRAS is highly expressed. Terminal positively charged regions increase the affinity of CPP-PNA conjugates for negatively charged DNA. Red represents peptide nucleic acid, blue represents cationic moieties, black represents the site of the G12D mutation in KRAS, minuses represent negative charge, pluses represent positive charge. **b,** AsPC-1 cells were treated with 500 nM FITC-CPP-PNA-G12D-7 for 4 hours prior to flow cytometry. Shown are the proportions of FITC+ gated cells, n = 3 per condition. **c-e,** Image quantification analyses (c-d) and representative fluorescent microscopy images (e) of AsPC-1 cells treated with 500 nM FITC-CPP-PNA-G12D-7 for 4 hours. n = 5 per condition. For all panels, statistical analysis was performed using two-tailed *t*-tests, **** p < 0.0001.

## Results

### PNA localization

The sequence regions of each PNA were designed using either a 17- or 20-mer sequence surrounding the region encoding the twelfth amino acid of KRAS G12D (Fig. 1a). As is standard in PNA design, base sequences were attached to a repeating peptide backbone, rather than the repeating phosphate backbone found in DNA. The regions surrounding the mutant KRAS sequence were wedged between a series of moieties featuring ionic and cationic regions flanked by hydrophobic regions, designed to improve cell penetration (Fig. 1a, Supplementary Fig. 1, Supplementary Table 1). All CPP-PNA conjugates were purified then validated by HPLC and matrix-assisted laser desorption/ionization mass spectrometry (MALDI MS) (Supplementary Fig. 2).

To quantify the ability of our modified PNAs to enter cells, we created FITC-CPP-PNA-G12D-7, a FITC-labeled version of CPP-PNA-G12D-7. We tested the ability to penetrate cells by flow cytometry in three lung and pancreatic cancer cell lines, AsPC-1, H358, and H1975. Attached cells were treated with 500 nM FITC-CPP-PNA-G12D-7 for 4 hours, followed by multiple washes to remove any remaining extracellular PNAs, and assessment for FITC by flow cytometry (Fig. 1b and Supplementary Fig. 3a-c). Over 99% of cells showed FITC signal, indicating that FITC-CPP-PNA-G12D-7 was highly competent at cell penetration.

We further assessed the subcellular localization of FITC-CPP-PNA-G12D-7 in the cytoplasm and nucleus by fluorescent microscopy. AsPC-1 cells dependent upon KRAS G12D and NCI-H358 cells dependent upon KRAS G12C were treated with FITC-CPP-PNA-G12D for 4 hours prior to staining for rhodamine-actin and DAPI. CPP-PNA conjugates were readily able to enter the nuclei of both G12D and G12C cell lines (Fig. 1c-e and Supplementary Fig. 3d-f). In fluorescence images, 74% of AsPC-1 cell nuclei had FITC foci, with an average of 1.6 foci of FITC signal per nucleus (Fig. 1c-e).

### Effects of G12D-PNA on binding, expression, and signaling

To evaluate the ability of CPP-PNA-G12D-7 to bind KRAS G12D DNA, we performed electrophoretic mobility shift assay (EMSA) using FAM-labeled complementary single-stranded or double-stranded DNA oligos. CPP-PNA-G12D-7 hybridized in a dose-dependent manner with the complementary KRAS G12D DNA sequence *in vitro*, with a 1:4 molar ratio proving sufficient to bind all ssDNA (Fig. 2a). At that dose, CPP-PNA-G12D-7 outcompeted the complementary sequence to reduce the amount of dsDNA (Fig. 2a). CPP-PNA-G12D-7 demonstrated selectivity by binding complementary G12D DNA at higher affinity than highly similar G12C DNA or WT DNA sequences (Supplementary Fig. 4).

**Figure 2.**
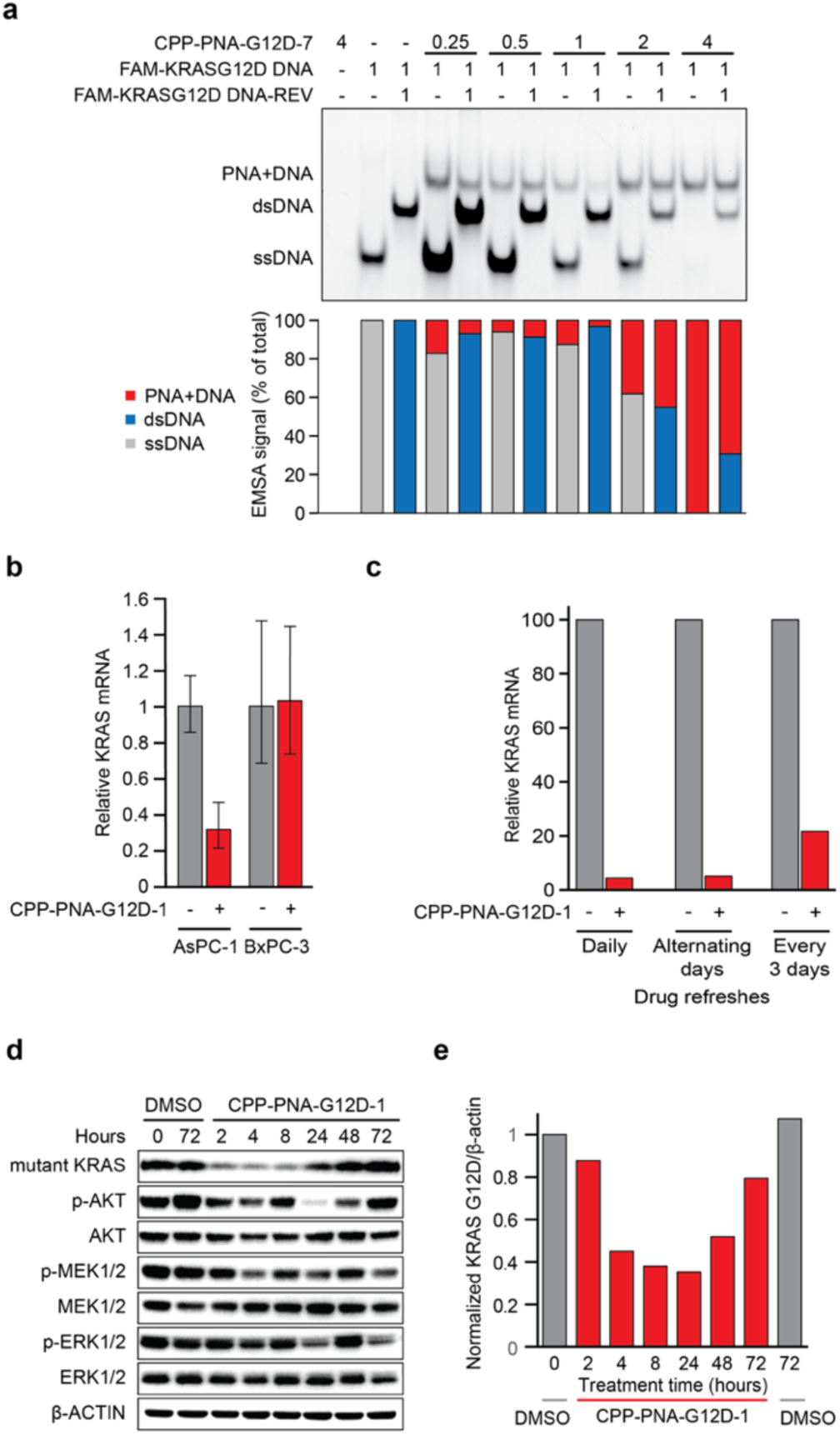
Expression of KRAS is repressed by CPP-PNAs. **a,** Electrophoretic mobility shift assay (EMSA) with CPP-PNA-7, complimentary FAM-labeled KRAS G12D DNA oligo, or reverse KRAS G12D oligo at the indicated molar ratios of 1 DNA to 0.25, 0.5, 1, 2, or 4 of PNA, with or without the reverse oligo. **b,** Quantitative real-time PCR from AsPC-1 and BxPC-3 cells treated with CPP-PNA-G12D-1 for 48 hours. **c,** Quantitative real-time PCR on AsPC-1 cells treated with CPP-PNA-G12D-1 for seven days using media changes daily, every 48 hours, or every 72 hours. **d-e,** Immunoblots (d), and image quantification (e), on lysates from AsPC-1 cells treated with CPP-PNA-G12D-1 or vehicle control for the indicated amount of time. Membranes were probed as indicated.

CPP-PNA conjugates can hybridize with complementary genomic DNA, resulting in a block of transcription for the target messenger RNAs.^21^ We treated AsPC-1 cells, which are homozygous KRAS G12D, and BxPC-3 cells, which are homozygous wild-type KRAS, for 48 hours with 500 nM of CPP-PNA-G12D-1. Expression of KRAS mRNA by qRT-PCR was repressed by about 70% in AsPC-1 cells, while no change was observed in BxPC-3, demonstrating selectivity of CPP-PNA- G12D-1 for cells expressing KRAS G12D (Fig. 2b).

To characterize the kinetics of KRAS mRNA depletion, we treated AsPC-1 cells for 7 days with CPP-PNA-G12D-1 and varied the length of time between when the cells were provided fresh media containing CPP-PNA-G12D-1. Daily and alternating day refreshments of CPP-PNA conjugate caused strong suppression of KRAS mRNA (Fig. 2c). The repression of KRAS mRNA was reduced when PNA refreshes were separated by 72 hours. Thus, PNAs retain their efficacy against mRNA for at least 48 hours.

Immunoblots using a KRAS G12D selective antibody demonstrated a drop in the amount of mutant KRAS beginning at 2 hours of G12D CPP-PNA treatment, culminating with a minimum at 8 hours, before a resurgence beginning at 24 hours and a return to pre-treatment amounts by 72 hours (Fig. 2d,e). CPP-PNA-G12D-1 inhibited signaling of multiple pathways downstream of KRAS over time. PNA treatment reduced p-AKT at most time points, with the effect lasting up to 72 hours (Fig. 2d and Supplementary Fig. 5a). CPP-PNA-G12D-1 slightly reduced p-MEK and p-ERK at later time points, lasting for 24 hours or more, followed by a rebound (Supplementary Figure 5b-c). These results led us to adopt a protocol of refreshing the CPP-PNA conjugate every 48 hours for subsequent cell viability experiments.

### CPP-PNA-G12D inhibition of viability of KRAS G12D-dependent cells

Selectively targeting mutant KRAS over wild-type KRAS is important for the ability to target cancer cells while sparing wild-type cells. To assess whether other KRAS G12D-dependent cell lines would be sensitive to inhibition with CPP-PNA-G12D-1 we treated a panel of six PDAC lines containing KRAS G12D and a variety of other secondary and tertiary driver mutations. In a dose-response experiment ranging from 62.5 - 2000 nM, CPP-PNA-G12D-1 suppressed viability in a dose-dependent manner across the panel of KRAS G12D mutant cell lines, with nanomolar IC50s (Fig. 3a).

**Figure 3.**
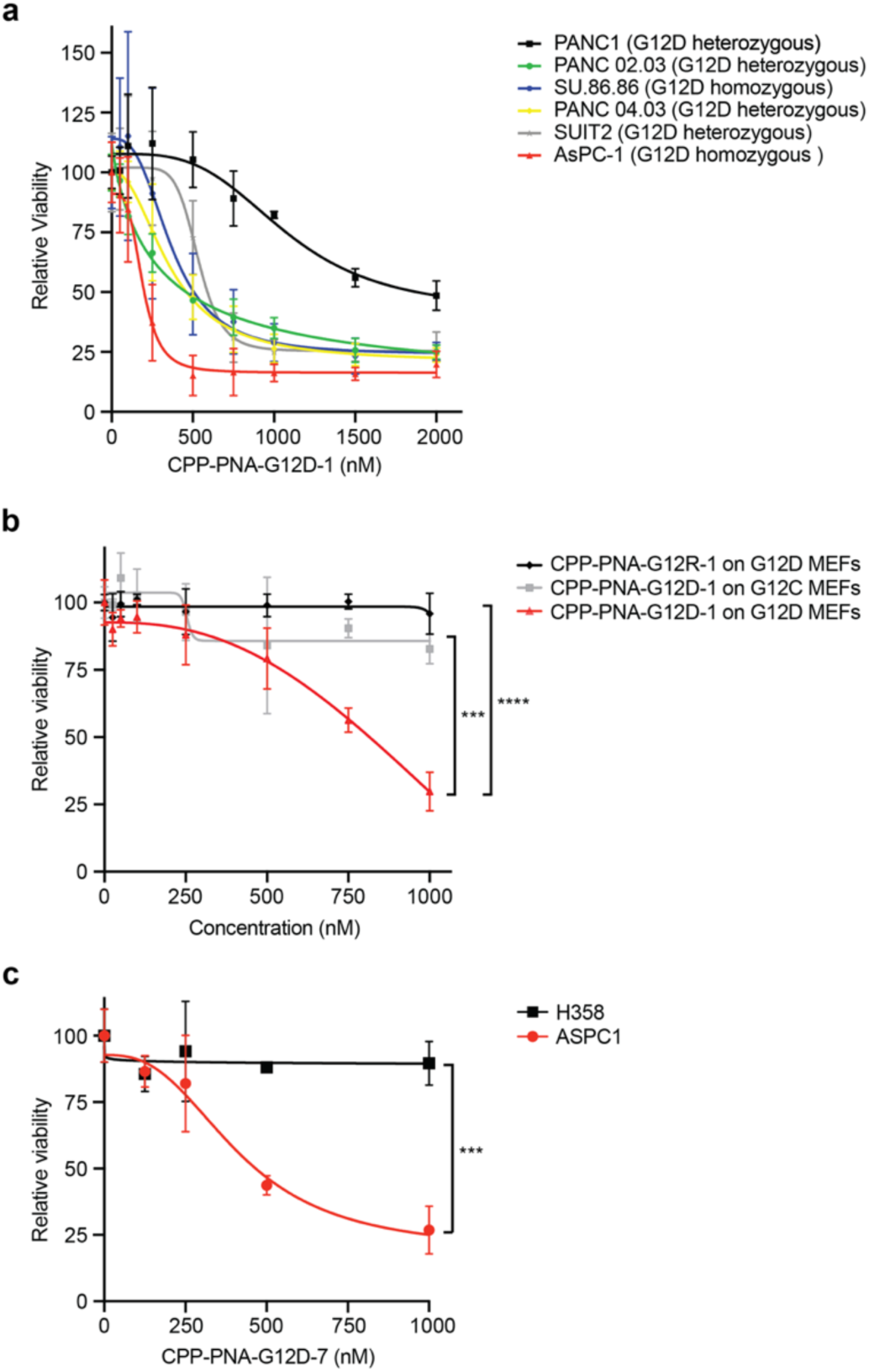
Efficacy and specificity of CPP-PNA-G12D against cell lines**. a,** Dose-response viability assays for six KRAS G12D PDAC cell lines following 7 days of treatment with media changes every 48 hours. **b,** Rasless MEFs containing no endogenous KRAS, NRAS, or HRAS that are overexpressing cDNAs with KRAS G12D (black and red lines) or KRAS G12C (grey line) following 7 days of treatment with CPP-PNA-G12D-1 or CPP-PNA-G12DR-1. **c,** Relative viability of sensitive AsPC-1 cells, dependent upon KRAS G12D, and resistant H358 cells, dependent upon KRAS G12C, following 7 days of treatment with the indicated doses of CPP-PNA-G12D-7. For all experiments, n = 3, *** p < 0.001, **** p < 0.0001.

To evaluate the specificity of our CPP-PNA-G12D conjugates, we employed isogenic MEF cell lines lacking endogenous KRAS, HRAS, and NRAS whose growth was rescued by cDNAs expressing either KRAS G12C or KRAS G12D.^22^ In 7-day dose course experiments, the CPP-PNA conjugate targeting G12D displayed a dose-dependent and genotype-specific effect in G12D MEFs (Fig. 3b). However, CPP-PNA-G12D-1 had no effect on negative control G12C MEFs, which only contain an off-target KRAS mutation. We developed a control CPP-PNA conjugates with the same backbone but with a nucleotide sequence targeting G12R called CPP-PNA-G12R-1 (Fig. 1a).Congruently, CPP-PNA-G12R-1 was incapable of inhibiting MEFs expressing KRAS G12D. These results, showing that mismatching the mutation in either the CPP-PNA or the target cells prevents inhibition, reinforced the selective nature of this CPP-PNA conjugate design and demonstrate single nucleotide level specificity of CPP-PNA-G12D-1 for KRAS G12D in cancer cells.

### Modifications to cell-penetrating peptide and length of KRAS binding sequence

Using our initial PNA design, we opted to iterate upon the sequence and structure toward increasing solubility and engagement with DNA. We employed several strategies to improve on our initial G12D PNA design including motifs to optimize specificity, cell penetration, and solubility. The KRAS G12D homologous base pair (bp) sequences were extended from 17 bp up to 20 bp to increase specificity, creating a genome-unique sequence. Alternative lengths, identities, and orders of the flanking region were designed and synthesized using longer lysine-rich amino acid flanking regions that were interspersed with histidines, arginines, or aspartic acids to make regions more or less cationic (Fig. 1a). CPP-PNA conjugates were designed with cell-penetrating regions concentrated in the N-terminal, the C-terminal, or both (Fig. 1a).

To evaluate the efficacy and specificity of the new PNA backbones and longer complementary sequences at targeting cells with KRAS G12D, we conducted 7-day relative dose course viability assays. These assays used KRAS G12D selective CPP-PNA conjugates with varying backbone content on either sensitive KRAS G12D AsPC-1 cells or insensitive KRAS G12C H358 cells. Each of these G12D-targeting CPP-PNA conjugates demonstrated significant specificity for inhibiting growth of AsPC-1 cells at higher concentrations (Fig. 3c and Supplementary Fig. 6). The most effective and selective version of our KRAS G12D PNA was CPP-PNA-G12D-7. This variant contained a 20 nucleotide KRAS G12D homologous region flanked on both ends by ε-palmitoyl-lysines. It encompassed aspartic acid and lysine-rich flanking regions on the 5’ end and a lysine-rich region on the 3’ end. (Supplementary Table 1) This candidate demonstrated an improved ability to inhibit cell viability in on-target cells, while sparing off-target cells (Fig. 4c).

**Figure 4.**
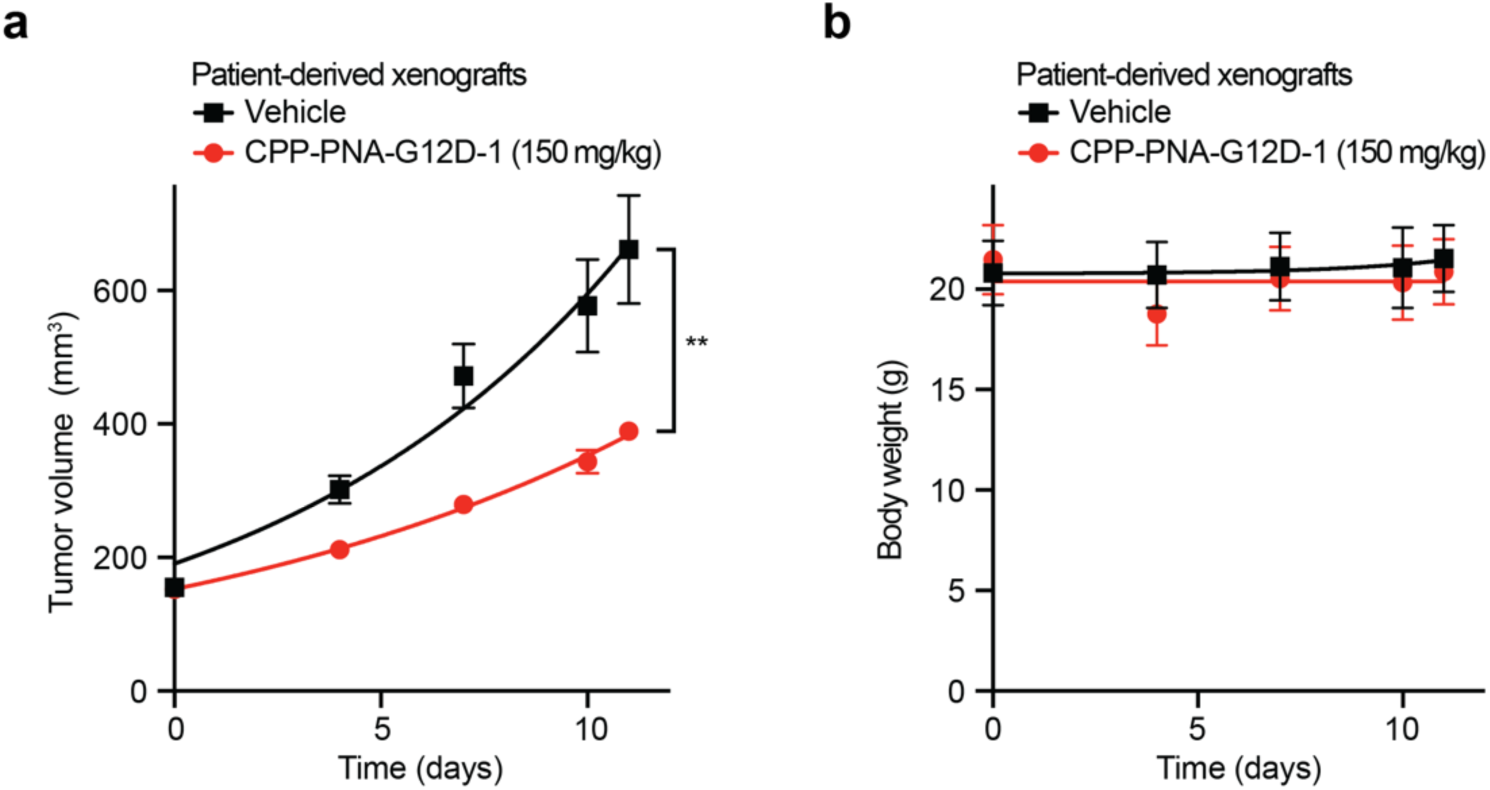
CPP-PNAs are effective and non-toxic in patient-derived xenografts of KRAS G12D PDAC. **a-b,** Tumor volume (a), and body weight (b), over time for mice bearing patient-derived PDAC xenografts treated with 150 mg/kg of CPP-PNA-G12D-1. Statistical analysis was performed using area under curve followed by *t-*test, n = 7, ** p < 0.001.

To confirm that the effect of our PNAs on cell viability is dependent on hybridization with cellular DNA, prior to treatment we pre-bound PNAs with single-stranded complementary DNA oligos to create heteromeric PNA/DNA oligo complexes. (Supplementary Fig. 7a,b) As expected, pre-bound PNA/DNA heteromeric oligo complexes were not effective at blocking cell viability in any genotype, including on-target G12D cells (Supplementary Fig. 7a,b). Thus, pre-binding of PNA to DNA prevented the PNA from being active against on-target cells, confirming that binding to complementary sequences is required for the PNA mechanism of action.

### Growth inhibition of KRAS G12D patient-derived xenografts in vivo

To evaluate the efficacy and safety of CPP-PNA-G12D *in vivo*, we employed a patient-derived xenograft (PDX) model derived from human PDAC. Mice treated with 150 mg/kg of CPP-PNA-G12D-1 demonstrated significantly less tumor growth relative to vehicle-treated controls (Fig. 4a). Upon dissection, some PNA had precipitated in the peritoneum, implying that only a fraction was absorbed. Treated mice did not show a significant change in body weight (Fig. 4b). CPP-PNA-G12D may represent an effective strategy for treating KRAS mutant cancers *in vivo*.

## Discussion

Cancer therapeutics currently available often do not definitively distinguish between normal cells and cancer cells, as they target pathways and signaling mechanisms found and used in both.^23–28^ Here, we created an improved delivery motif conjugated to complementary PNAs, making them reliable transcription suppressors. By extending and rearranging the cationic delivery peptide motif, we generated PNA-peptides which suppress transcription with sufficient sensitivity to distinguish single-point mutations.^16^ These modifications were applied to a PNA oligomer-delivery peptide conjugate to suppress gene transcription within expressed regions. The PNA oligomer differentiated sequences differing by a single nucleotide base, demonstrating clear specificity for KRAS G12D over other KRAS sequences. These results support the efficacy and specificity of this improved PNA-delivery peptide conjugate strategy. Findings were validated in a tumor-bearing xenograft resulting in inhibition of cell proliferation. Improved solubility for *in vivo* systems would be expected to increase efficacy in mice. By design, our PNAs are resistant to intra and extracellular enzymes, show single base-pair mismatch discrimination, when conjugated to transport/nuclear-locating peptides are permeable to nuclear and cell membranes, and specifically suppress genes at nanomolar concentrations in cell culture. These oligonucleotide analogues lend themselves well to automated peptide synthesis incorporating stabilizing peptides, which make these oligomers considerably more membrane-permeable and more conducive to helical binding.

The present study focused on KRAS G12D as it is a prevalent cancer-causing mutation not targeted by clinically approved KRAS inhibitors sotorasib and adagrasib. Our PNAs differ from earlier KRAS PNA designs in several significant ways. Previous KRAS targeting PNAs were not able to effectively penetrate cell membranes without lipofectamine or 2D-layered Mg-Al layered double hydroxides (LDHs) as carriers for delivery.^18,19^ This is a significant limitation, as lipofectamine is not generally suitable for clinical applications.^18^ Due to the ionic and cationic modifications added to the ends of the PNAs used in this study, they were capable of penetrating the cell membrane and localizing within the nucleus of KRAS mutant cells. Prior PNAs were designed to target the KRAS mRNA through complementarity with the forward sequence, whereas our approach focuses on targeting the mutation exclusively at the DNA level through complementarity with the reverse strand.^18,19^ Due to the statistical challenge for the PNA complement finding its chromosomal target sequence, the cationic termini of the CPP-PNA oligomer are stabilized against the polyanionic chromosomal target better ensuring that the PNA conjugate remains within the chromosomal binding arena and improving initiation of duplex binding. In addition, the CPP-PNA conjugates in the present study demonstrated a greater reduction in targeted cell viability at lower concentrations.^18^

In contrast to many prior strategies, such as chemotherapy, most monoclonal antibodies, and most small molecule inhibitors, PNA strategies target tumor-promoting gene sequences that are inherent only to cancer cells and incorporate several therapeutic advantages. Experimental pan-Ras inhibitors such as RMC-6236, currently in clinical trials, are capable of binding wild-type KRAS and also inhibit several off-targets such as NRAS and HRAS present in normal cells.^8^ By design, obstructing transcription through targeting the gene sequence of a mutation/translocation site is unaffected by secondary mutations elsewhere within the gene. In contrast, inhibitors that rely upon the specificity of a protein binding site may be affected by mutation-driven structural perturbations occurring anywhere within the entire protein structure, leading to therapeutic resistance, as observed in patient samples treated with G12C-selective KRAS inhibitors adagrasib or sotorasib.^8,29,30^ If the target sequence mutates or a new dominant subtype arises, it can be sequenced in order to amend the complementary PNA sequence much more rapidly than the development of a revised small molecule inhibitor. Unlike siRNA-mediated suppression strategies, PNA-delivery peptide conjugates do not require reliance upon cellular mechanisms such as RISC processing, which may predispose siRNAs to off-target suppression. PNAs are less susceptible than RNAi to nucleases or other enzymatic degradation. They may be chemically modified, enabling versatile functional derivatizations. While CRISPR-Cas9 systems are powerful gene editing tools with remarkable clinical uses, there are limitations for employing CRISPR against cancer.^31^ Although CRISPR has uses *ex vivo*, the *in vivo* clinical utility of CRISPR-based strategies presents obstacles such as the reliance on viral delivery, the daunting prospect of incorporating every cell with bacterial nucleases and possible constitutive expression, and permanent genetic modifications.

The ability to target cancer cells specifically through their genetic distinctiveness presents a more specifically effective approach toward treatment of cancer. Using PNAs to reversibly obstruct the transcription or replication of a gene sequence, such as an oncogene, that is both crucial for tumorigenesis and specific to the cancer cell itself, offers a potential route to facilitate such an approach. These results indicate that PNAs show promise as both a tool for studying point-mutation-driven tumors and as a possible therapeutic strategy capable of sparing normal cells while selectively targeting mutant cancer cells.

Since PNA strategies depend upon targeting a complementary gene sequence, they can in principle be applied to suppress the expression of any oncogenic mutation or other disease-causing genes for laboratory studies or potential clinical development. Several early successes have been reported. The oncogene Myc was silenced using a γPNA-based platform designed to target genomic DNA *in vivo*.^32^ Sites were selected upstream of key transcription start points to block transcription factor binding, effectively inhibiting oncogene expression.^33^ Transcriptional inhibition of MYC resulted in the downregulation of genes involved in DNA replication and repair, making lymphoma cells more responsive to chemotherapy.^32,33^ Peptide nucleic acids that downregulate MDM2 activate p53 to enhance tumor suppressor activity.^34^ An MDM2 PNA conjugated to 9-aminoacridine was effectively delivered to cells with and without a carrier and targeted a cryptic AUG initiation site on *MDM2* mRNA. The PNAs reduced MDM2 protein levels and increased p53 transcription and protein levels, resulting in reduced cell viability.^34^ The PNA OLP-1002 is being evaluated for its therapeutic potential in a range of chronic pain conditions, including osteoarthritis, diabetic neuropathic pain, trigeminal neuralgia, and chemotherapy-induced pain.^35^ OLP-1002 is an SCN9A antisense PNA that selectively inhibits the Nav1.7 sodium channel. In a Phase 2a trial, a single subcutaneous injection of OLP-1002 significantly reduced neuropathic hypersensitization and provided long-lasting therapeutic effects. PNAs in combination with conventional antibiotics have been used to reduce bacterial resistance, focusing on targeting resistance genes in strains such as *H. pylori, A. baumannii, P. aeruginosa, H. influenzae,* and more.^36–40^ Since PNAs cannot easily penetrate bacterial membranes, they were conjugated with the permeable agent (KFF)3K to enhance delivery for Gram-negative bacteria.^40^ PNA conjugates exhibited strong antibacterial activity against both Gram-negative and Gram-positive bacteria, with effectiveness dependent on factors including PNA length, binding site, and conjugant type.^40^ In addition, PNAs are being explored as antiviral agents as their high binding affinity and resistance to degradation make them effective for targeting specific viral RNA sequences, such as in Hepatitis B virus and the SARS coronavirus.^41,42^

PNAs have been employed in neurological settings to target the CNS mu opioid and neurotensin receptors in rats to study their role in neurotensin responses.^43^ The PNAs were designed to bind to the mRNA or DNA of these receptors. They found that PNA treatment reduced the neurotensin-induced pain relief and temperature changes, with a decrease in NT receptor binding sites in the periaqueductal gray and hypothalamus, while sparing control rats. Moreover, PNAs are also being tested in neurodegenerative diseases including Huntington’s disease and Alzheimer’s disease by targeting and modulating gene expression associated with these conditions.^13,44,45^ In addition, PNAs have shown promise for genetic diseases like Duchenne Muscular Dystrophy and cystic fibrosis.^46–48^ These studies suggest that PNAs may cross the blood-brain barrier, supporting their use for inhibiting protein expression in the brain.^43,49^

PNAs are useful for detection and imaging of diseases. KRAS PNAs have been used as reagents for peptide nucleic acid-locked nucleic acid polymerase chain reaction (PNA-LNA PCR) to detect KRAS mutations in surgically resected lung cancer samples, with more efficient detection than traditional PCR methods.^50^ PNAs can detect other biomarkers, including EGFR and lncRNAs associated with prognostic outcomes.^51,52^ In addition to cancer, PNAs have been used as radiolabeled biomarkers for tracking bacterial and fungal infections.^53,54^

Cell-penetrating PNAs present an exciting intervention for selectively targeting cancer-driving genes such as KRAS. Our findings that CPP-PNA-G12D-1 inhibited on-target KRAS G12D transcription in pancreatic ductal adenocarcinoma cell lines and patient-derived xenografts provide proof-of-principle for the efficacy and selectivity of the programmable and modular cell-penetrating peptide conjugated PNA technology.

## Methods

### Cell Lines

AsPC-1 (RRID:CVCL_0152), NCI-H1975 (RRID:CVCL_1511), NCI-H358 (RRID:CVCL_1559), SU.86.86 (RRID:CVCL_3881), SUIT-2 (RRID:CVCL_3172) cells were maintained in RPMI-1640 (Gibco #11875119), 10% fetal bovine serum (FBS) (R&D Systems #S11550H), and 1% Penicillin/Streptomycin/L-Glutamine (PSG) (Gibco #10378016) as their complete media. PANC.02.03 (RRID:CVCL_1633) and PANC.04.03 (RRID:CVCL_1636) cells were maintained in RPMI-1640 (Gibco #11875119), 10% FBS, and 1% PSG, with the addition of 20 U/ml human recombinant insulin (Sigma-Aldrich #I-1882) to create their complete media. PANC-1 (RRID:CVCL_0480) cells were maintained in DMEM (Gibco #11965092), 10% FBS, and 1% PSG as their complete media. Rasless MEFs expressing KRAS G12C or KRAS G12D (NCI RAS Initiative/Leidos Biomedical Sciences) were cultured and maintained in DMEM (Gibco #11965092), 10% FBS, and 1% PSG, with the addition of 4 ug/L Blasticidin S (Thermo Scientific Chemicals #J61883FPL) to create their complete media. All cells were maintained at 5% CO_2_ in a 37°C humidified incubator. Cells used for assays were counted using a Countess 3 Automated Cell Counter (Invitrogen #AMQAX2000) and plated accordingly. All cell line identities were confirmed using short tandem repeat analysis (ATCC #135-XV). All cell lines tested negative for mycoplasma using MycoAlert® Mycoplasma Detection Kit (Lonza #LT07-318).

### PNA production

PNAs were designed in collaboration with Oncogenuity, Inc. PNAs were produced by PNA Bio. Modifications to PNA oligomers include acetylation (Ac), palmitoylation (palm), or FITC conjugation (see Supplementary Table 1). Underlined letters represent nucleotides. High performance liquid chromatography was performed using an Agilent 1100 Series from Agilent Technologies. Matrix-assisted laser desorption/ionization mass spectrometry was performed using an AXIMA-Assurance from Shimadzu Biotech.

### Viability assays

For dose-course experiments, live cells were plated at 5,000 cells/well in sterile optical-bottom 96-well plates (Thermo Scientific #165306) in their respective complete media on day 1. To reduce the impact of edge effects, triplicates were spatially separated on the plate and edge wells were not read. On day 2, each PNA was diluted in media to desired concentrations, then cells were treated with the indicated concentration of media with PNA or vehicle, with all wells receiving the same dose of vehicle. AsPC-1, NCI-H358 cells were treated for 72 hours with no drug refresh or for 7 days with drug refreshes every 48 hours. At the experimental endpoint, brightfield microscopy images were taken with EVOS FL Auto Imaging system (Applied Biosystems #AMEX- 1200). Relative viability was then assessed using CellTiter-Glo® Luminescent Cell Viability Assay (Promega #G7572) per manufacturer protocol and placed in Varioskan Lux (Thermo Scientific #VLBLATGD2) or a Spectramax i3 (VWR #10192-220). Data were normalized to vehicle-treated cells within cell lines.

### Fluorescent microscopy

Approximately 50,000 AsPC-1 and NCI-H358 live cells were plated on 12-well glass bottom plates (Mattek Corp #P12G-1.5-10F). On the following day, cells were treated with either DMSO or 500 nM FITC-labeled PNA (FITC-CPP-PNA-G12D-7) for 4 hours. The cells were fixed using 4% formaldehyde (Thermo Scientific Chemicals #J60401-AP). After 15 minutes at room temperature, the formaldehyde was removed and the wells were washed with PBS. The cells were permeabilized using 0.1% Triton X-100 in PBS (Fisher Chemicals #BP151-500). After 10 minutes at room temperature, the wells were washed with PBS. Rockland blocking buffer (Rockland Immunochemicals #MB070) was added for 30 minutes. Wells were washed with PBS. Invitrogen ActinRed™ 555 ReadyProbes™ Reagent Rhodamine phalloidin (Sigma-Aldrich #R37112) was mixed with 1% BSA + PBST (Fisher BioReagents #BP9703100) and placed in the wells to stain for actin. The plates were incubated on an orbital shaker overnight away from light. The next day, cells were washed with PBS, then mounting media containing DAPI (Vector Laboratories #H120010) was added to stain for DNA. For imaging, a Nikon A1 Confocal Microscope with NIS-Elements software was used. Five image fields of each well were captured with 60X objective oil immersion. CellProfiler was used to quantify nuclei, dots representing PNAs, and actin, and the same settings were applied to all images, n = 5 per condition.

### Mouse xenografts

Female BALB/c nude mice 6-7 weeks of age were injected with patient-derived xenografts (PDXs) from pancreatic ductal adenocarcinomas (PDACs), n = 3 per condition. Tumors were treated on alternating days with 150 mg/kg of CPP-PNA-G12D-1 or vehicle for two weeks. Mice were weighed at each time point. Upon sacrifice, tumor, liver, heart, lung, pancreas, and kidneys were collected and analyzed.

### Immunoblots

AsPC-1 cells were treated with either DMSO or 800 nM of CPP-PNA-G12D-1 for the indicated number of hours. Cells were harvested, lysates were quantified using BSA, and lysates were run on Criterion TGX Pre-cast gels (Bio-rad), transferred to PVDF membranes, then were probed with the following antibodies from Cell Signaling Technologies: KRAS G12D (Cat #14429), pERK1/2 (Cat #4370), ERK1/2 (Cat #4695), p-MEK1/2 (Cat #9154), MEK1/2, p-AKT (Cat #4060), AKT (Cat #2920). Membranes were imaged using an Odyssey Imager (LI-COR).

### Quantitative PCR

AsPC-1 cells were treated for 7 days with media changes every 24 hours, 48 hours, or 72 hours. RNA was extracted using an RNeasy Mini Kit (Qiagen) and screened for quality and concentration using a Nanodrop spectrophotometer. Reverse transcription was performed using High Capacity cDNA Reverse Transcription Kit (Applied Biosystems Inc) following manufacturer instructions. Quantitative PCR was performed in 384-well format using a Fast PCR System 7900H (Applied Biosystems Inc). qPCR primer sequences are listed in Supplementary Table 3. dCt values were calculated against GAPDH and ddCt were calculated against vehicle-treated controls.

### Flow Cytometry

Approximately 50,000 AsPC-1, NCI-H1975, and NCI-H358 cells were incubated with either FITC- CPP-PNA-G12D-7 or negative control DMSO for 4 hours. After the incubation, cells were trypsinized, neutralized with normal media, centrifuged at 300 x g for 5 min at 4°C, aspirated, resuspended in flow media, and added to a 96-well U-bottom plate (Sigma-Aldrich #CLS4515). Flow cytometry was performed on a CytoFLEX S flow cytometer (Beckman Coulter #C09762). Live cells were gated with side-scatter and forward-scatter (Fig. 1b and Supplementary Fig. 3a,b). FITC-CPP-PNA-G12D-7 cells were gated using parameters in which unstained controls had ≤ 0.2% positive cells using Flowing Software 2.5.1.

### Electrophoretic Mobility Shift Assay (EMSA)

Lyophilized KRAS WT, KRAS G12C, and KRAS G12D DNA oligomers were synthesized by Sigma-Aldrich, where each forward sequence were fluorescein-labelled (FAM) and their corresponding reverse complement sequences were not FAM-labelled (Supplementary Table 2).

DNA oligomers were reconstituted in 1X PBS. 0.06828 nmol of the indicated DNA oligo, with or without the indicated molar ratio of PNA, were heated to 90°C, then gradually cooled to anneal as described in Thadke et al.^55^ Each sample was loaded onto a 10% non-denaturing PAGE in 1X TBE buffer (Thermo Scientific #B52) and then separated at 120V for 60 minutes. Gel images were processed and visualized by the ChemiDoc (Bio-Rad #12003154) via UV-Transilluminator.

### Statistical Analysis

Statistical tests are described in figure legends. All analyses were performed in GraphPad Prism 10 (RRID:SCR_002798) or Microsoft Excel (RRID:SCR_016137). For dose-course experiments, data were processed using the area under the curve (AUC); following the processing, p-values were calculated using one-way analysis of variance (ANOVA) followed by Tukey’s multiple comparisons test in GraphPad Prism. Mouse xenograft data were processed using the area under the curve (AUC); following the processing, p-values were calculated using an unpaired *t-*test in GraphPad Prism. Gated cell proportions and fluorescence data were compared using two-tailed *t*-tests in GraphPad Prism. From the above statistical analyses, p-values are shown as: ns represents not significant, * p < 0.05, ** p < 0.01, *** p < 0.001, and **** p < 0.0001.

## Acknowledgments

The authors gratefully acknowledge Leonard J. Ash and Nikita Meghani for insight on PNAs, Jill Bargonetti for use of the ChemiDoc MP Imaging System, Lloyd Williams from the Bio-Imaging Facility of Hunter College for use of the Nikon A1 Confocal Microscope at Belfer Research Building, Joon Kim from the Flow Cytometry/Genomic Lab of Hunter College for use of the Becton-Dickinson BD CytoFLEX S flow cytometer, and all members of the Wolfe lab for insightful advice. PNA Bio synthesized CPP-PNA conjugates. Rasless MEF cells were obtained from the National Cancer Institute Ras Initiative/Leidos Biomedical Research. We thank Ottavia Busia-Bourdain, Diana P. Bratu, Paul Feinstein, and Elesha McGrath and for providing feedback on the figures and manuscript.

