## Supplementary Figures for "Selective targeting of oncogenic KRAS G12D using peptide nucleic acid oligomers attached to cell-penetrating peptides"

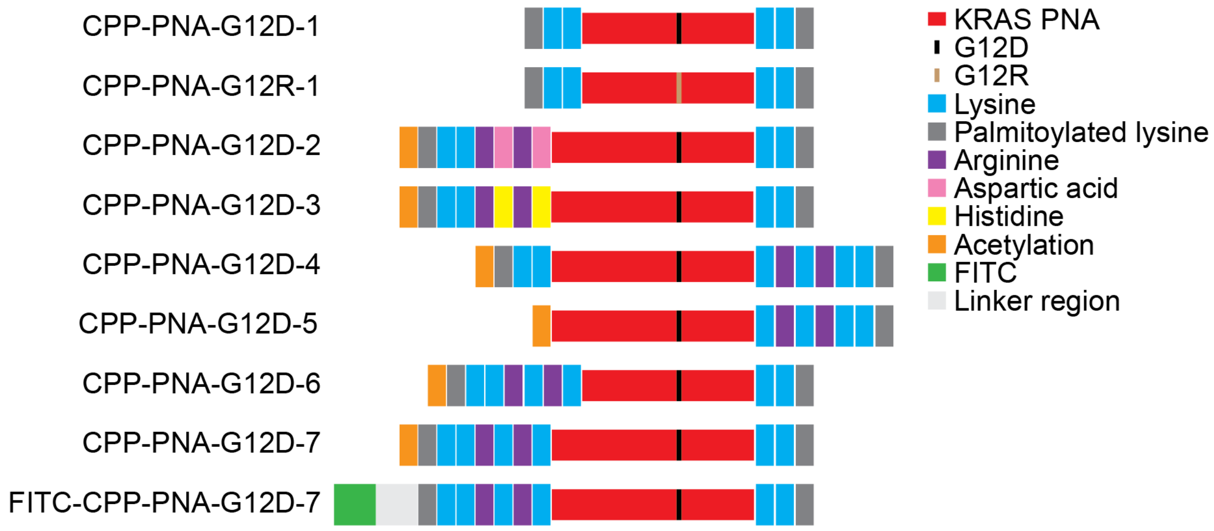


**Supplementary Figure 1.** Diagram of PNA structures. Red= complementary KRAS PNA sequence, black = G12D, light brown = G12R, blue = lysine, dark grey = palmitoylated lysine, purple = aspartic acid, pink = arginine, yellow = histidine, orange = acetylation, green = fluorescein isothiocyanate (FITC), light grey = linker region.

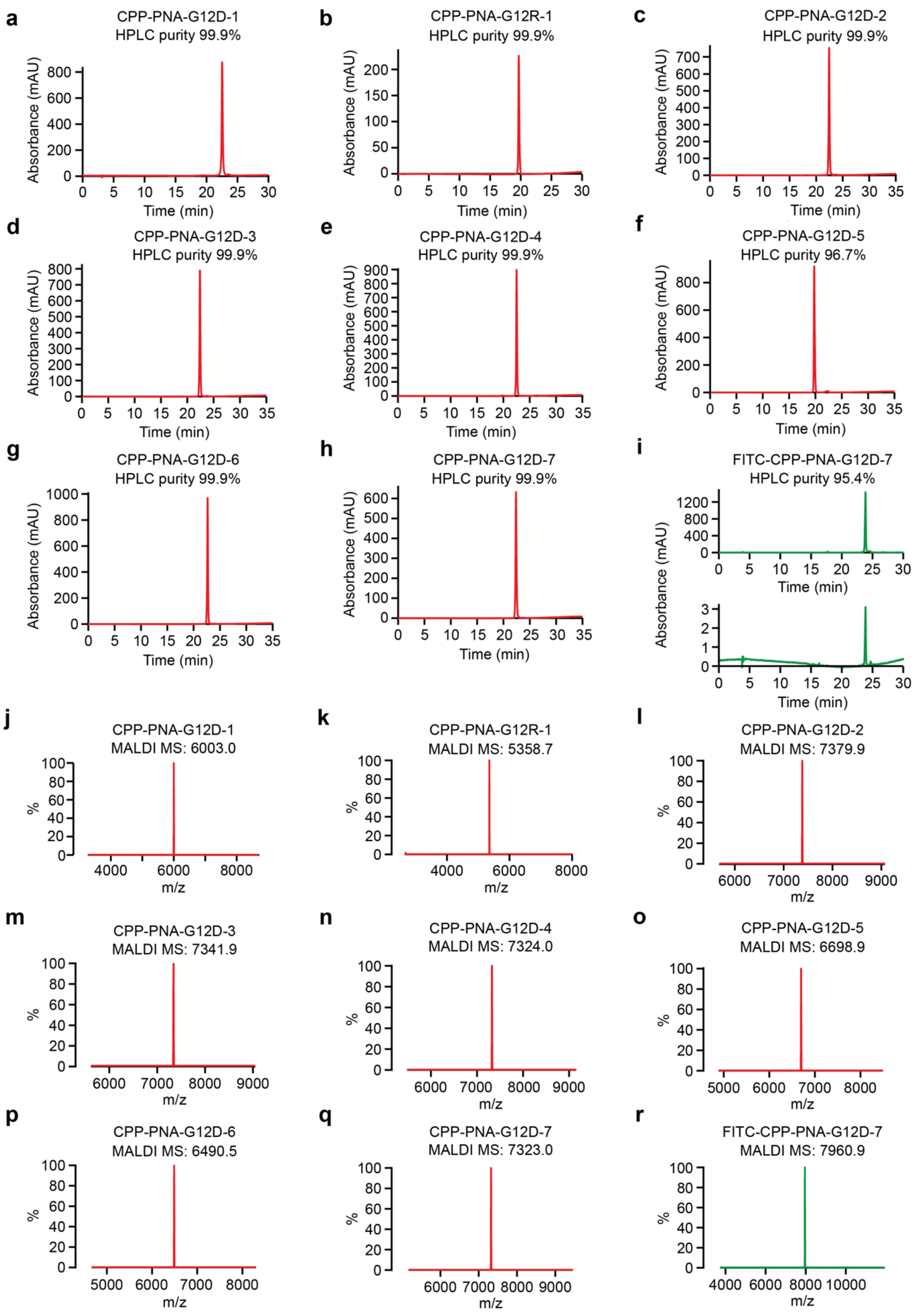
a
CPP-PNA-G12D-1
HPLC purity 99.9%
Absorbance (mAU)
Time (min)
b
CPP-PNA-G12R-1
HPLC purity 99.9%
Absorbance (mAU)
Time (min)
c
CPP-PNA-G12D-2
HPLC purity 99.9%
Absorbance (mAU)
Time (min)
d
CPP-PNA-G12D-3
HPLC purity 99.9%
Absorbance (mAU)
Time (min)
e
CPP-PNA-G12D-4
HPLC purity 99.9%
Absorbance (mAU)
Time (min)
f
CPP-PNA-G12D-5
HPLC purity 96.7%
Absorbance (mAU)
Time (min)
g
CPP-PNA-G12D-6
HPLC purity 99.9%
Absorbance (mAU)
Time (min)
h
CPP-PNA-G12D-7
HPLC purity 99.9%
Absorbance (mAU)
Time (min)
i
FITC-CPP-PNA-G12D-7
HPLC purity 95.4%
Absorbance (mAU)
Time (min)
Time (min)
j
CPP-PNA-G12D-1
MALDI MS: 6003.0
%
m/z
k
CPP-PNA-G12R-1
MALDI MS: 5358.7
%
m/z
l
CPP-PNA-G12D-2
MALDI MS: 7379.9
%
m/z
m
CPP-PNA-G12D-3
MALDI MS: 7341.9
%
m/z
n
CPP-PNA-G12D-4
MALDI MS: 7324.0
%
m/z
o
CPP-PNA-G12D-5
MALDI MS: 6698.9
%
m/z
p
CPP-PNA-G12D-6
MALDI MS: 6490.5
%
m/z
q
CPP-PNA-G12D-7
MALDI MS: 7323.0
%
m/z
r
FITC-CPP-PNA-G12D-7
MALDI MS: 7960.9
%
m/z

**Supplementary Figure 2.** Validation of CPP-PNA conjugates. **a-i,** Purity of each CPP-PNA was assessed by high-performance liquid chromatography (HPLC) for CPP-PNA-G12D-1 (a), CPP-PNA-G12R-1 (b), CPP-PNA-G12D-2 (c), CPP-PNA-G12D-3 (d), CPP-PNA-G12D-4 (e), CPP-PNA-G12D-5 (f), CPP-PNA-G12D-6 (g), CPP-PNA-G12D-7 (h), FITC-CPP-PNA-G12D-7 (i). **j-r,** Purity of each CPP-PNA was assessed by matrix-assisted laser desorption/ionization mass spectrometry (MALDI MS) for CPP-PNA-G12D-1 (j), CPP-PNA-G12R-1 (k), CPP-PNA-G12D-2 (l), CPP-PNA-G12D-3 (m), CPP-PNA-G12D-4 (n), CPP-PNA-G12D-5 (o), CPP-PNA-G12D-6 (p), CPP-PNA-G12D-7 (q), FITC-CPP-PNA-G12D-7 (r). Indicated peak values are MS found (M+1).

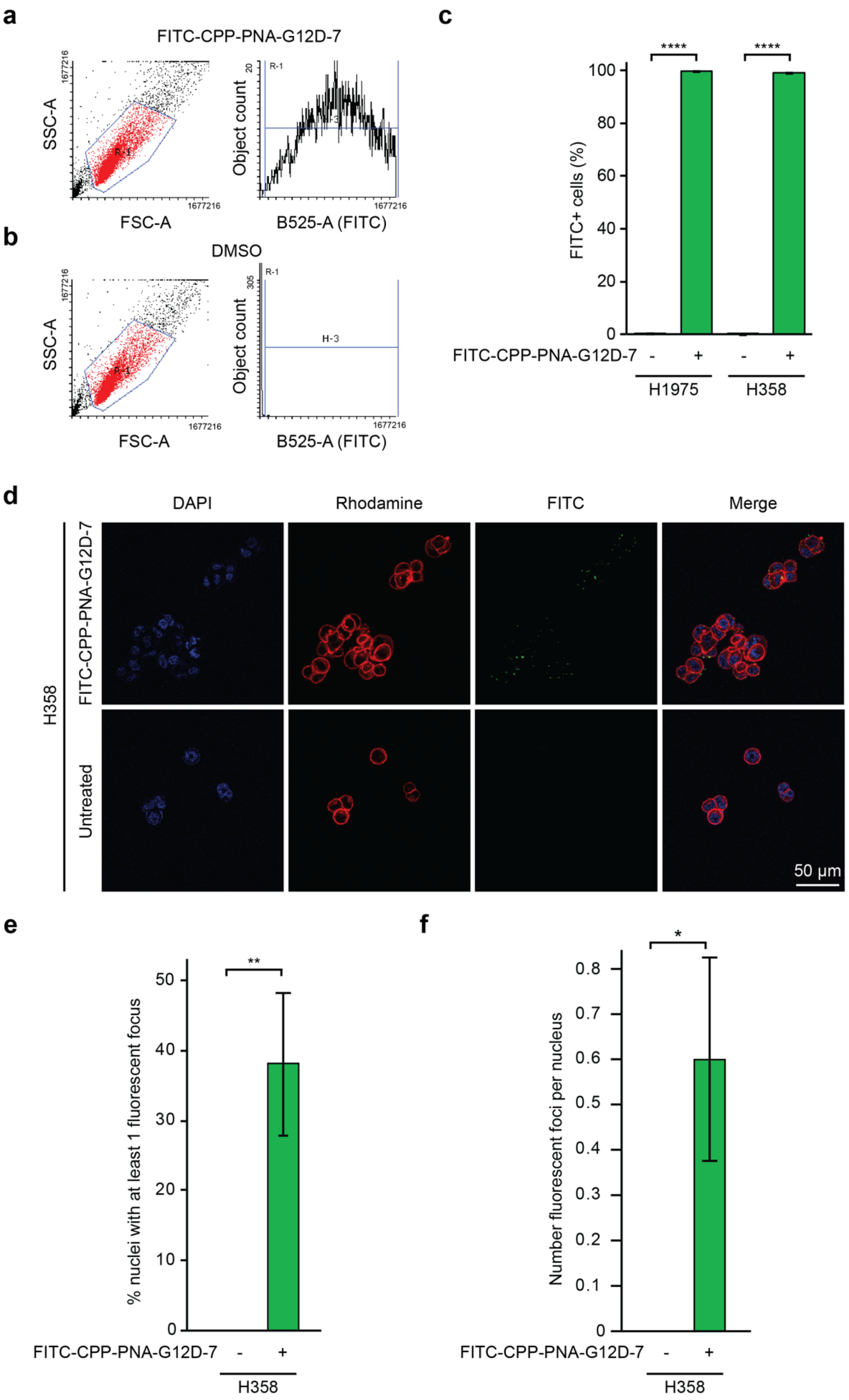
a
FITC-CPP-PNA-G12D-7
SSC-A
FSC-A
Object count
B525-A (FITC)
b
DMSO
SSC-A
FSC-A
Object count
B525-A (FITC)
c
FITC+ cells (%)
FITC-CPP-PNA-G12D-7
H1975
H358
d
DAPI
Rhodamine
FITC
Merge
H358
FITC-CPP-PNA-G12D-7
Untreated
50 μm
e
% nuclei with at least 1 fluorescent focus
FITC-CPP-PNA-G12D-7
H358
f
Number fluorescent foci per nucleus
FITC-CPP-PNA-G12D-7
H358

**Supplementary Figure 3.** Cell penetration of PNAs in additional cell lines. **a-b,** Representative flow cytometry gating and results for AsPC-1 cells treated with FITC-CPP-PNA-G12D-7 (a) or DMSO (b) for 4 hours prior to flow cytometry, n=3. **c,** H1975 and H358 cells were treated with FITC-CPP-PNA-G12D-7 for 4 hours prior to flow cytometry. Shown are the proportions of FITC+ gated cells, n = 3 per condition. **d-f,** Representative fluorescent images (d), and image quantification analyses (e-f), of H358 cells treated with FITC-CPP-PNA-G12D-7 for 4 hours. n = 5 per condition. For all panels, statistical analysis was performed using two-tailed *t*-tests, * $p < 0.05$, ** $p < 0.01$.

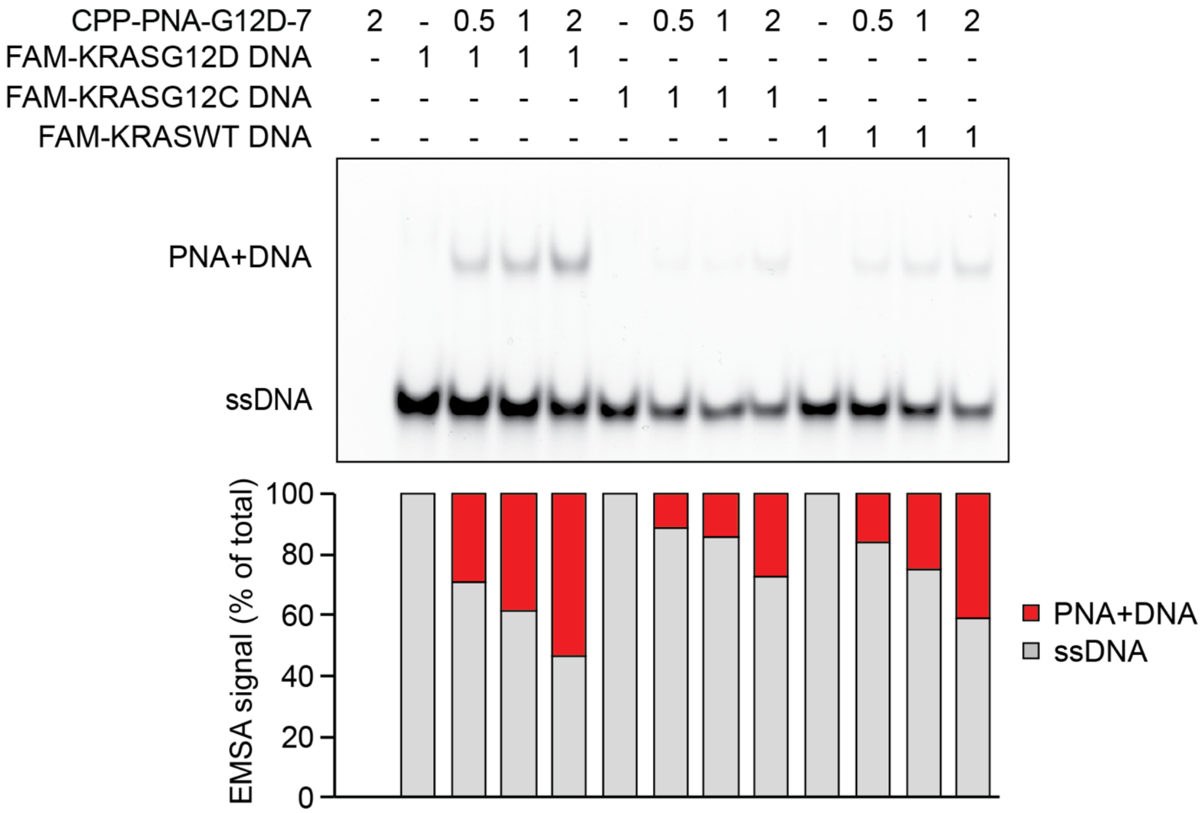


**Supplementary Figure 4.** EMSA using FAM-labeled DNA oligos of the indicated genotype incubated with CPP-PNA-G12D-7 at 1 DNA:0.5 PNA, the equimolar ratio of 1 DNA:1 PNA, or 1 DNA:2 PNA.

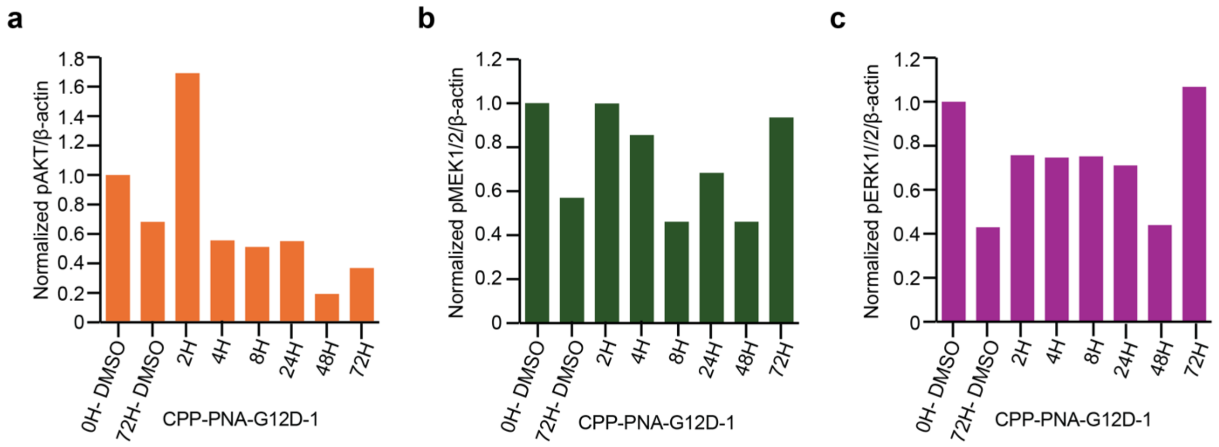


**Supplementary Figure 5.** Quantification of time-course signaling. **a-c,** Quantification of time-course immunoblot images lysates treated with DMSO or CPP-PNA-G12D-1 of **a,** pMEK1/2, **b,** pERK1/2, and **c,** pAKT. Each quantification is normalized to β-actin.

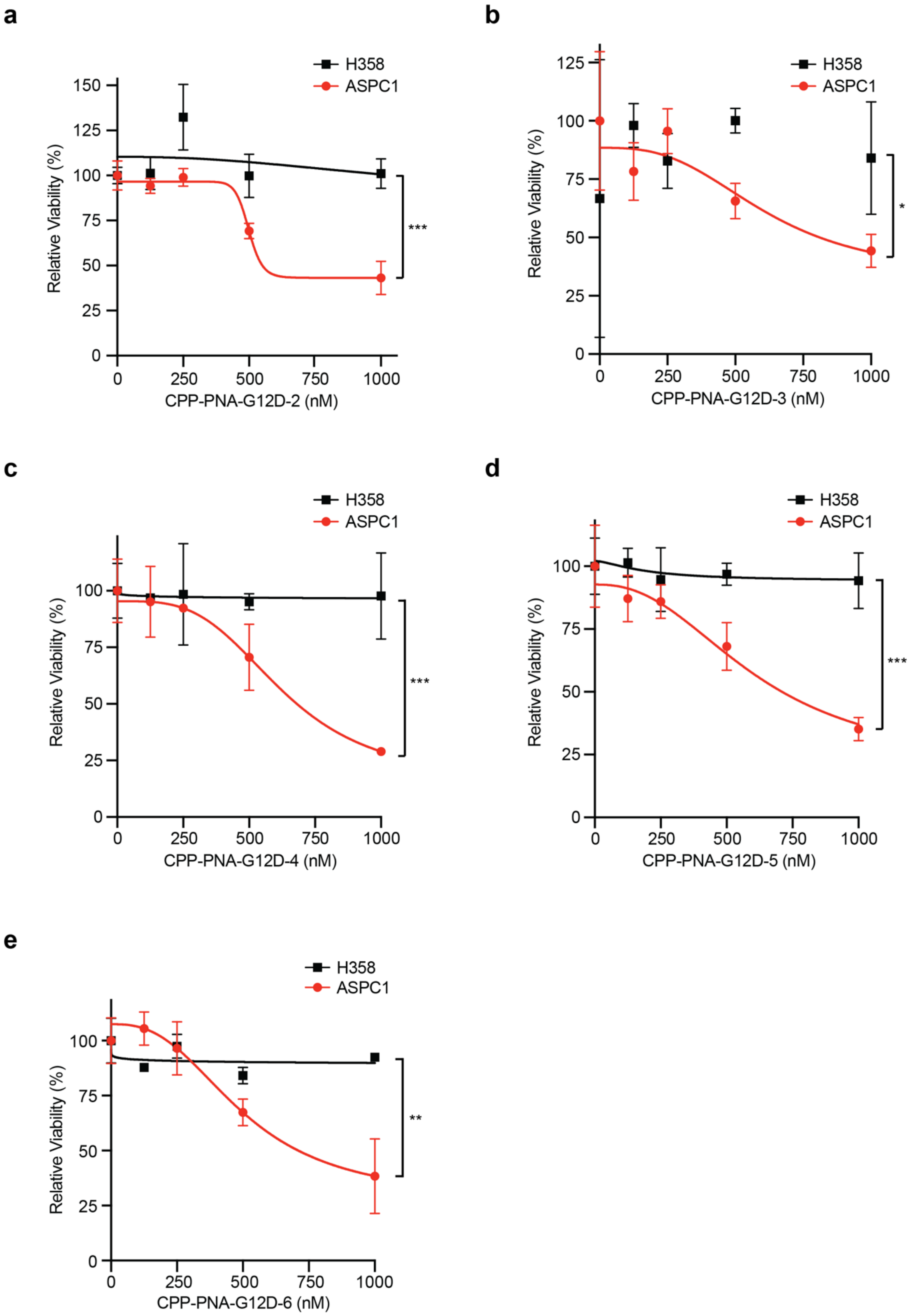
a
H358
ASPC1
Relative Viability (%)
150
125
100
75
50
25
0
0
250
500
750
1000
CPP-PNA-G12D-2 (nM)
***
b
H358
ASPC1
Relative Viability (%)
125
100
75
50
25
0
0
250
500
750
1000
CPP-PNA-G12D-3 (nM)
*
c
H358
ASPC1
Relative Viability (%)
100
75
50
25
0
0
250
500
750
1000
CPP-PNA-G12D-4 (nM)
***
d
H358
ASPC1
Relative Viability (%)
100
75
50
25
0
0
250
500
750
1000
CPP-PNA-G12D-5 (nM)
***
e
H358
ASPC1
Relative Viability (%)
100
75
50
25
0
0
250
500
750
1000
CPP-PNA-G12D-6 (nM)
**

**Supplementary Figure 6.** PNAs selectively target KRAS G12D mutants. **a-e,** Relative viability of sensitive AsPC-1 cells, dependent upon KRAS G12D, and resistant H358 cells, dependent upon KRAS G12C, after 7 days of treatment with the indicated doses of CPP-PNA-G12D-2 (a), CPP-PNA-G12D-3 (b), CPP-PNA-G12D-4 (c), CPP-PNA-G12D-5 (d), or CPP-PNA-G12D-6 (e). See Supplementary Fig. 1 and Supplementary Table 1 for more information on these CPP-PNA conjugates. For all panels, statistical analysis was performed using area under the curve (AUC), $n = 3$, ** $p < 0.01$, *** $p < 0.001$.

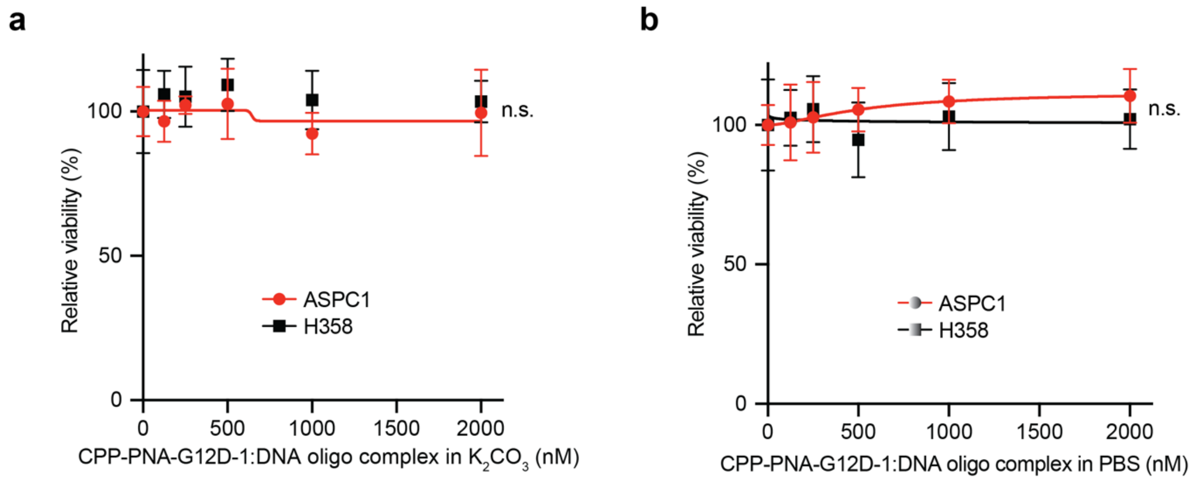


**Supplementary Figure 7.** Pre-binding of PNA with DNA oligo complexes impairs efficacy. **a,** Pre-incubation of CPP-PNA-G12D-1 with complementary DNA strands prevented efficacy against AsPC-1 cells (red) or H358 cells (black) at all tested doses, irrespective of whether the PNA was dissolved in $K_2CO_3$ (a) or PBS (b) as a buffer.
